# The occasional perils of reflection (across the midline; in geometric morphometrics)

**DOI:** 10.1101/770677

**Authors:** David C. Katz

## Abstract

Manually collecting landmark data on a large biological sample takes a long time. Several options exist to speed data collection, though each strategy introduces problems or raises concerns of its own. For bilaterally symmetric structures (e.g., crania), some recent papers recommend limiting landmark collection to one side and the midline, then “mirror-reflecting” landmarks across the midline to produce an approximation of the true bilateral configuration. However, where the midline is narrow relative to the bilateral anatomy, net midline landmark deviations from the mid-sagittal axis or plane will distort the mirror-reflected configuration. Here, I test whether this is a substantive concern at the scale of real biology. To do so, I simulate small amounts of mediolateral error on the mean shape from a sample of human mandibles (*n* = 178), then compare the distribution of simulated forms to variation in the data. I also test how faithfully mirror-reflected configurations replicate bilateral shape and size relationships. In both analyses, midline deviations from symmetry create striking distortions. I go on to show that incorporating a small number of landmarks from the opposite side of the mandible produces far more accurate estimates of bilateral shape than does mirror reflection. Mirror reflection is clearly inappropriate for these data and is likely suspect in all cases of narrow midline morphology.

## Overview

Since its emergence in the late 1980s (1–3), the number of biological questions assessed using geometric morphometrics (GM) has continued to expand (4, 5). As the scope of these studies has become more ambitious, sample size requirements often increase as well. Classification with canonical variates analysis calls for several times more cases than coordinate dimensions (6). Adequately quantifying within- and between-group shape variation in a widespread and/or ecologically diverse species or across a species-rich taxonomy can require many hundreds or even thousands of observations (7–9), as can identifying the genetic underpinnings of shape and form (10, 11).

Large samples pit scientific objectives against time and funding constraints. Efficiencies are needed. One can speed data collection by reducing the number of landmarks. However, sparser landmarks sets are less able to capture local changes. Morphometricians are advised to landmark “sufficiently” (12, p. 25), a necessarily vague term. In the absence of reliable criteria for determining the optimum number of landmarks to include, “most geometric morphometric studies are characterized by an oversampling of (anatomical) landmarks as an exploratory strategy: it allows for unexpected findings (and nice visualizations).” (13) Recent efforts to quantify sufficiency (14) implicitly support the inference that more landmarks are better. Landmarking can be completed more quickly if the work is divided among co-investigators (15), or even crowd-sourced (8), but these introduce additional sources of error. Automated and nearly automated landmarking allow for rapid phenotyping of shape and form (16–19). However, automated approaches are still in relatively early stages of deployment. Moreover, the 3D digital samples (CT, laser scan, etc.) required for automated landmarking are themselves time-consuming to collect and process (20), and not yet available for many specimen collections.

For user-placed landmarks on bilaterally symmetric features, an additional option is to sample only one side. In an idealized system, bilaterally symmetric features are anatomical pairs subject to identical developmental regimes, resulting in left and right versions which are mirror images of one another. Bilateral symmetry is far from the only kind of symmetry to have emerged over the course of organismal evolution, but it is the most common kind (21, 22). It is often subdivided into matching and object symmetry subcategories. Matching symmetric features are those that have physically distinct right and left versions (femora, innominates, wings, etc.). Object symmetric features are those whose bilaterally symmetric right and left sides are joined at an anatomical midline to form a single object (cranium, articulated pelvis, sphenoid, etc.) (23, 24).

Where asymmetries between right and left are not an important aspect of the investigation, bilateral symmetry can be leveraged for faster data collection by making the right or left side the subject of analysis. This practice is referred to herein as a “focal side” approach. If a specimen’s focal side is damaged or missing, an investigator can often landmark and then reflect the opposite side to incorporate it into the sample (25). For object symmetric structures, focal side data collection entails collecting landmarks on one side and along the midline (together, a hemi-form). If the better preserved side is incomplete, it is often possible to record the antimeres and impute these landmarks to the focal side by reflected relabeling (26). These strategies can cut the time needed for GM data collection nearly in half, and thus nearly double *n* under a given time or financial constraint.

However, Generalized Procrustes Analysis (GPA, 27) can be unkind to hemi-form data. In GM, GPA (described further in Materials & Methods below) is the most widely used algorithm for superimposing landmark configurations in a common coordinate space, and prerequisite to the ordinations and statistical analyses typically conducted for shape data. For object symmetric structures, bilaterally placed landmarks tend to balance the configuration during superimposition, such that the midline landmarks lie fairly close to a plane (Fig. 1B). If asymmetries are removed from the data (24, 26), the resulting midlines are, in fact, planar (Fig. 1A). In contrast, with a focal side approach, midline variation in the hemi-forms is no longer structured (or balanced) by the similarity of the right and left sides. This can result in somewhat larger, oblique differences among the midlines of the observations (Fig. 1C).

**Fig 1.**
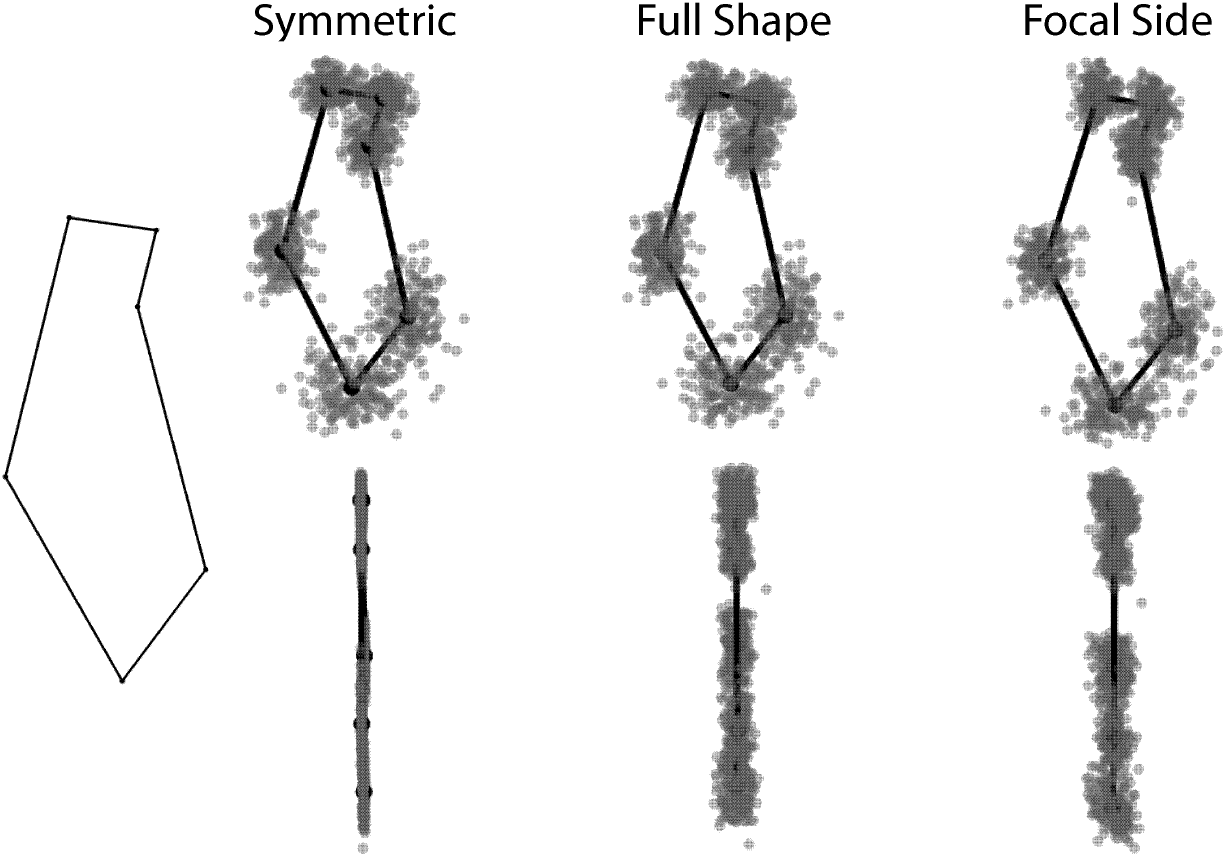
Medial and anterior views of the midline symphysis landmarks from symmetric, whole shape, and focal side superimpositions of human mandibular configurations (*n* = 178). Landmark map provided in Fig. 4.

Each of the superimpositions in Figure 1 is technically correct. However, the results of focal side superimposition can be difficult to interpret, at least visually. Though Procrustes superimposition aligns configurations statistically, not biologically, it is preferable and reassuring to work with arrangements that more closely accord with our intuitions about how biology varies (28). Hemi-form registrations of object symmetric structures sometimes want for this intuitive quality.

More importantly, as Cardini recently demonstrated through a series of comparisons (29, 30), bilateral and focal side superimpositions do not paint identical pictures of overall similarities and differences among subjects in a sample. This raises concerns about whether the efficiencies of the focal side approach are sufficient to justify its use.

Cardini’s proposed solution is “mirror reflection” of hemi-forms (29–31). Mirror reflection is a special case of reflected-relabeling in which only the midline landmarks are used to fit a hemi-form to its reflection (because midlines are the only landmarks a hemi-form and its reflection have in common). The end result is a bilaterally symmetric configuration with a linear (2D landmarks) or planar (3D) midline (Figure 2).

**Fig 2.**
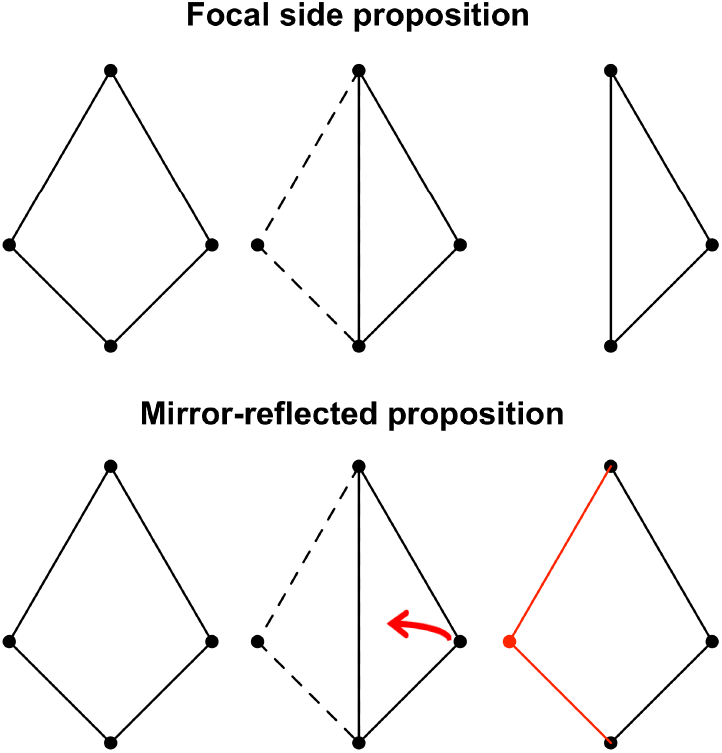
Schematic of focal side and mirror reflection procedures. Left polygons: true configurations. Middle polygons: in both cases, landmarks are collected on one side and along the midline. Right polygons: the focal side approach assesses the resulting hemi-form; mirror reflection fits the hemi-form to its reflection along the midline, creating a bilaterally symmetric form.

Cardini evaluated the efficacy of mirror reflection in bilateral landmarks datasets consisting of insect exoskeletons and vertebrate crania from eleven taxa. For each dataset, he considered whether focal side or mirror-reflected configurations better replicated the Procrustes distance and centroid size relationships observed for the full complement of bilateral landmarks. For most datasets (those considered 30), he expands the comparison to include the symmetric component of bilateral shape variation (24), as well. For all comparisons, symmetric component shape relationships are consistently the most similar to bilateral shape relationships. This finding is no surprise, as the symmetric and bilateral configurations differ only by asymmetry, the magnitude of which is often small and/or non-directional. In all but one comparison, bilateral configuration shape relationships were better approximated by mirror-reflected data than by hemi-form data. The consistently higher correlations for mirrored data led Cardini to recommend that investigators consider mirror reflection as an efficient data collection strategy.

However, Cardini’s samples all share a characteristic that makes mirror-reflection more likely to produce good (more similar to bilateral) results: the landmarks along the midsagittal define a midline that is deep relative to the other dimensions of the landmark configuration. Why should this matter? Recall that with mirror reflection, a bilateral shape is constructed by superimposing the recorded midline to its reflection. The position of the reflected landmarks is thus dependent on the extent of (net) deviation of the midline landmarks from an idealized midsagittal plane.

Figure 3 illustrates the relationships between midline depth, asymmetry in midline landmarks, and mirror-reflected shape distortion. The top row presents the “true” positions of four landmarks on three two-dimensional configurations. The configurations are identical except for the y-axis location of the inferior midline landmark. The red arrows indicate identical landmark placement errors (though anatomical deviations from symmetry that are confined to the midline would produce the same effect). The contrast between mirror-reflected reconstructions and true shapes is shown in the bottom row.

**Fig 3.**
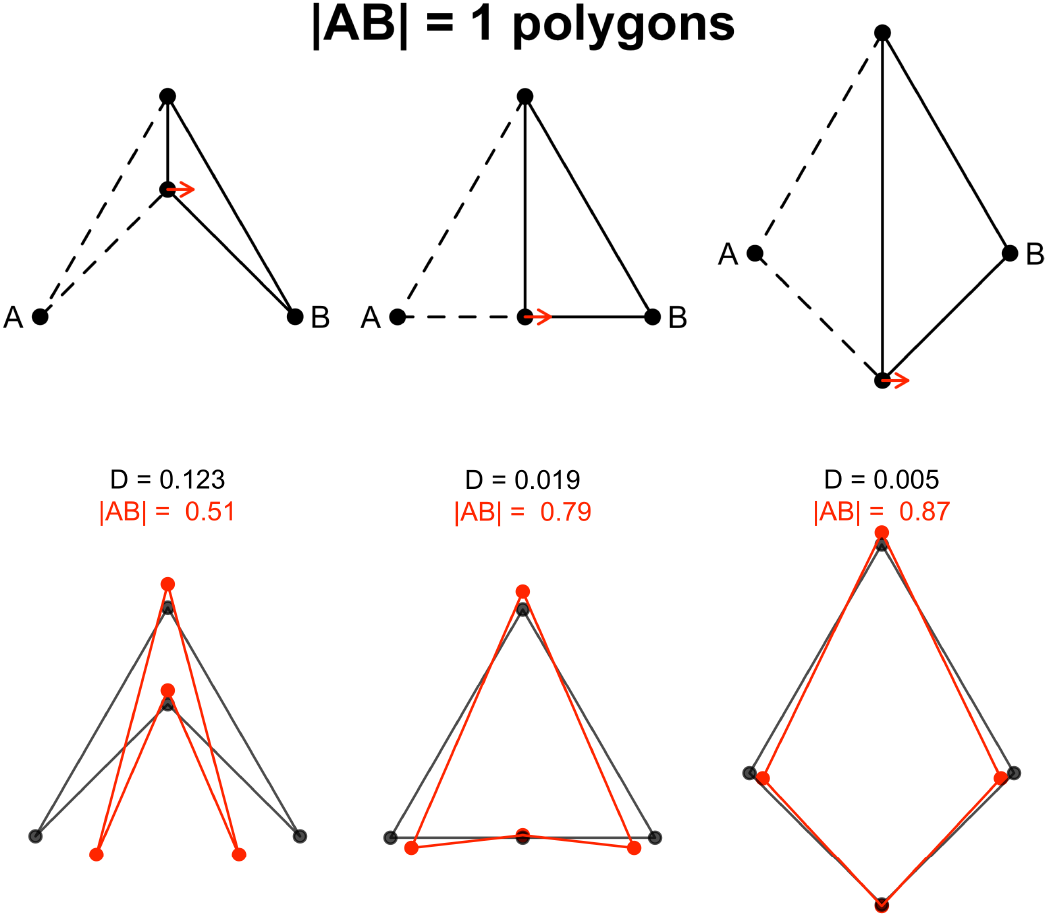
*Top row:* Three configurations with |AB| = 1 vary only in the vertical position of the inferior midline landmark. Identical mediolateral errors (red arrows) of 0.1 units are committed when recording the right side for mirror reflection. *Bottom row:* The Procrustes distances (D) between true and mirrored shapes, and the difference in |AB|, are most severely affected for the narrow midline configuration.

The extent to which midline error distorts true shape is clearly a function of the depth of the midline axis. When the midline is antero-posteriorly deep (right column of Figure 3), modest deviations still allow for reasonable mirror approximations of bilateral shape. When the midline is narrow relative to the bilateral configuration (left column of Figure 3), midline asymmetry is propagated out to bilateral landmarks. Small deviations are liable to produce large shape and size differences between mirror-reflected and true bilateral shapes.

Moreover, the distorting effect of midline deviations are magnified with increasing distance from the symphysis much like, for a given angular rotation, the distance traveled along the circumference of a circle increases as function of the length of the radius. The situation is analogous to how choice of baseline can produce a spectrum of outcomes with Bookstein’s two-point registration (32). Typically, the narrower the baseline, the more baseline variance influences interlandmark correlations and the results of statistical analysis (12 pp. 57-59, 33).

The most obvious vertebrate anatomy for which mirror reflection would be suspect is the mandible (at least for species with a fused symphysis). Mandibles are likely second only to crania –the landmark-rich darlings of GM—as shape subjects (Figure 4). The issue, then, is a matter of general concern for GM practitioners.

**Fig 4.**
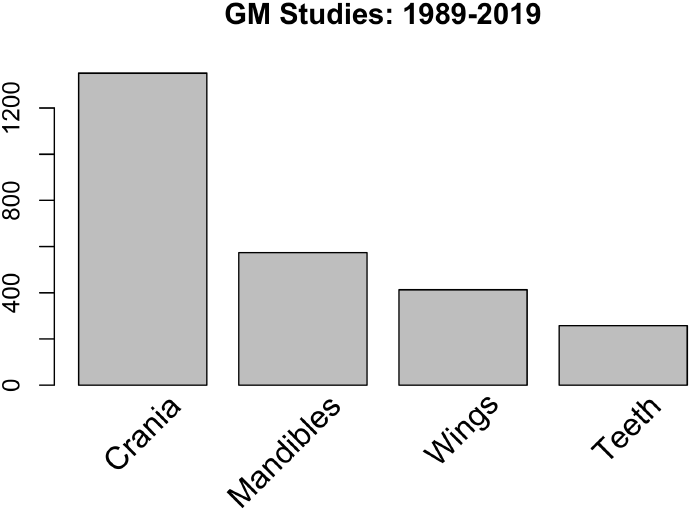
The four most prominent anatomical features studied with GM, based on an informal topic search in Web of Science (https://apps.webofknowledge.com), using the bar label terms in the figure and similar terms for the same anatomy.

Below, I use a sample of human mandibles to quantify how severely mirror-reflection affects results of shape analysis. I simulate small, lateral deviations at the midline landmarks of the symmetric mean for the full complement of landmarks, then create mirror reflections from the right hemi-forms. Through various comparisons of the simulations to the bilateral observations, I demonstrate that for narrow-midline morphology, even small midline deviations produce severe distortions. Comparison of shape and size relationships among observations using alternative landmarking strategies raise the same concerns. Finally, I consider the implications of these findings, and suggest an alternative solution that is nearly as efficient as the focal side approach but far better at recovering bilateral shape.

## Materials and Methods

All analysis was performed in R (34), primarily with custom routines. GPA and symmetric GPA were performed using functions from geomorph (35). For some routines, the shapes package (36) was used for Ordinary Procrustes Analysis (superimposition of two configurations). Figure 1 was generated in rgl (37) and captured by screenshot.

### Data

The mandibular data are adult, bilateral, complete cases (*n*=178) from the larger, global sample of human mandibles described in (7). The landmark data consists of six paired, bilateral landmarks that capture the mandibular condyle and lateral tooth row. Depending on the analysis, the configurations include either two (arch subset) or six (full configuration) midline landmarks on the mandibular symphysis (Figure 5; Table 1).

**Table 1.** Landmarks, starred (*) if bilateral.

| ID | Name | Description |
| --- | --- | --- |
| id | infradentale | Labial alveolar midline septum between central incisors |
| mo | mandibular orale | Lingual symphyseal midline between central incisors |
| im | incurvatio mandibularis | Maximum posterior ingress of anterior symphysis |
| me | mental eminence | Anteriormost projection of mandibular symphysis |
| gn | gnathion | Inferiormost projection of mandibular symphysis |
| ms | mental spine | Symphyseal midline superior to termination of mental spines |
| i2* | Incisor margin | Lingual alveolar margin between lateral incisor and canine |
| m2* | lingual M2 | Lingual alveolar margin in coronal plane with lingual groove of M2 |
| pm* | posterior molar | Most posterior point in the tooth row, relative to a coronal plane |
| m1* | buccal M1 | Buccal alveolar margin in coronal plane with buccal groove of M1 |
| cp* | C-P3 margin | Labial alveolar margin between canine and P3 |
| cd* | mandibular condyle | Most posterior point on mandibular condyle |

**Fig 5.**
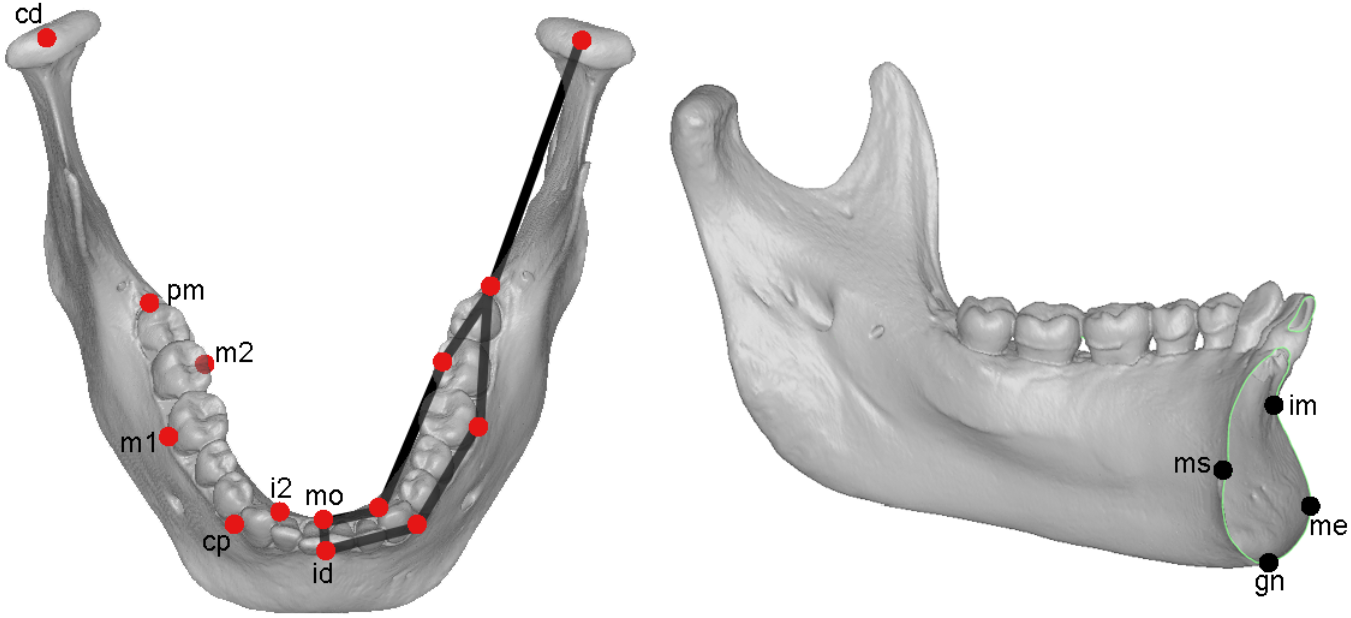
Landmark map. Red landmarks: 14-landmark arch subset. Red + black landmarks: 18-landmark full configuration.

### Simulations on observations

The distortion introduced by small deviations at the midline upon mirror reflection is evaluated with simulations. All simulations follow the same basic procedure. First, the original landmark configurations are superimposed *via* symmetric GPA. In the GPA procedure, configurations are centered at the origin and scaled to unit centroid size, after which each is rotated so as to minimize its summed squared distance from an iterated estimate of the mean shape (38). The remaining differences among configurations are considered shape differences. Centroid size, the inverse of the scaling factor from the superimposition, provides an estimate of specimen size. The symmetric version of GPA (24) requires expansion of the sample to include both the original configurations and their reflections. After superimposition of this expanded sample, each configuration and its reflection are averaged, yielding a symmetric shape for analysis.

I use the mean forms (mean shape scaled to mean size) of the arch subset and full configuration superimpositions as templates to generate 1000 simulated arch subset and full configuration forms. Simulating on forms instead of shapes is useful for generating simulated data that varies at a biologically meaningful scale. Prior to simulation, in order to ensure that deviations occur only in a mediolateral direction, the means were rotated so that anatomical axes and Cartesian axes were the same. I then simulated uncorrelated deviations at the midline landmarks along the Cartesian axis that was coincident with the mediolateral direction.

Deviations—one for each midline landmark—were sampled at random from a normal distribution, *N*(0; 0.25 mm). With a standard deviation of 0.25 mm, each simulated deviation is expected to be well within the range of generally accepted measurement error (1–2 mm) in osteometry (39). Note that though the standard deviation for the random sampling exercise is guided by a measurement error standard, for purposes of the analyses, it is irrelevant whether the midline landmark deviations from an idealized midsagittal are thought of as error or off-center biology. Finally, I create a reflected version of the simulated right hemi-form, and join the two at the midline to create a mirror-reflected configuration. The midlines of the simulated hemi-shape and its reflection are fit using Ordinary Procrustes Analysis (OPA), which minimizes the squared distance between a reference and target (38). The effect of midline deviations on shape and size are evaluated both visually and statistically.

### Shape Recovery

I test the performance of an alternative strategy to mirror reflection that also approximates bilateral mandibular shape without using the full complement of landmarks. This alternative is referred to herein as shape recovery. Relative to mirror reflection and focal side approaches, it expands data collection to include a small number of landmarks from the opposite side. Creating a bilateral configuration then becomes a standard reflected relabeling task: create a reflection of the configuration, swap right and left landmark locations in the data matrix, rotate one to the other by OPA of the landmarks they share in common, then average common landmarks (27). As with mirror reflection, what remains is a symmetric configuration.

I test two shape recovery hypotheses:

- *H_1_. For narrow midline configurations, incorporation of opposite side landmarks will recover bilateral shape better than does mirror reflection*. Once opposite side landmarks are included, the relationship of the sides to one another is largely determined by the position of the opposite landmarks relative to their antimeres on the focal side. The effect of midline deviations from symmetry become confined to the symphysis itself. In contrast, with mirror reflection, the relationship of the sides to each other is entirely a function of net deviations at the midline.
- *H_2_: Bilateral shape is better recovered by incorporation of opposite side landmarks located more distant from the midline*. As with the effect of midline deviations on bilateral landmarks, deviations from symmetry between opposite side landmarks and their antimeres become magnified at more distal landmarks (if any). Thus, the opposite side landmarks located furthest from the midline should better recover bilateral shape because there will be no magnification of deviations.

I test these hypotheses by attempting to recover bilateral shape with three opposite side landmark pairs: proximal (*ii* and *cp*), middle (*m1* and *m2*), and distal (*pm* and *cd*). See Figure 5 for landmark locations. I then follow and extend the approach of Cardini (29, 30) to compare the shape and size relationships for the true configurations and their derivative approximations (mirror-reflected, shape recovery, symmetric, and focal side). Right hemi-shapes are used for the focal side, mirror-reflected, and shape recovery reconstructions. For each dataset, I compute matrices of pairwise Procrustes and centroid size differences among observations. I then compute the pairwise correlations among these matrices in order to evaluate whether any of the strategies that use less than the full complement of landmarks capably replicates the structure of variation in complete forms.

Both hypotheses have a sample-wide focus; for any given observation, mirror reflection may outperform shape recovery. Moreover, the generality of the results presented below are subject to variation among bilateral landmarks in measurement error and asymmetry, which I do not evaluate here, and which may differ among species and samples.

## Results

### Simulations

I simulated mediolateral deviations at the midline toothrow landmarks of the arch subset symmetric mean 1000 times, each time mirror-reflecting the right side to generate 1000 simulated versions. While many of the simulated outcomes (Figure 6) are similar to the true configuration (e.g., within the 50% quantile range), very large distortions are common, too.

**Fig 6.**
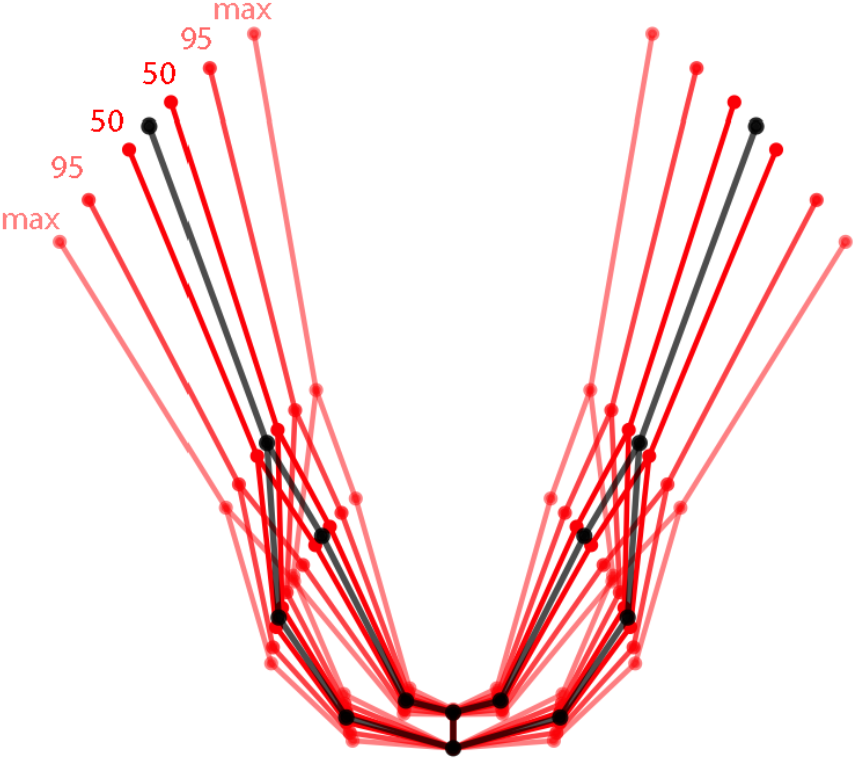
Arch shapes produced by simulating mediolateral deviations—*N*(0; 0.25 mm)—at the two midline landmarks of the mean arch subset hemi-form (black wireframe), then mirror-reflecting the right side. Successively faded red wireframes indicate 50% and 95% quantiles and the maxima of the range of simulations on the mean. Quantiles derived based on intercondylar width. For purposes of this illustration, the plotted configurations were translated such that the all anterior midline landmarks are positioned at the origin.

Figure **7A** projects the arch subset mirror-reflection simulations into the morphospace of the first two principal components (PCs) of the superimposed sample observations. Many of the simulations lie well outside the range of human variation in the sample. Figure 7B displays the corresponding simulations and data for the full configuration. While distortions are reduced relative to the arch subset, the range of variation for simulated outcomes still spans most of the range of sample variation on PC1.

**Fig 7.**
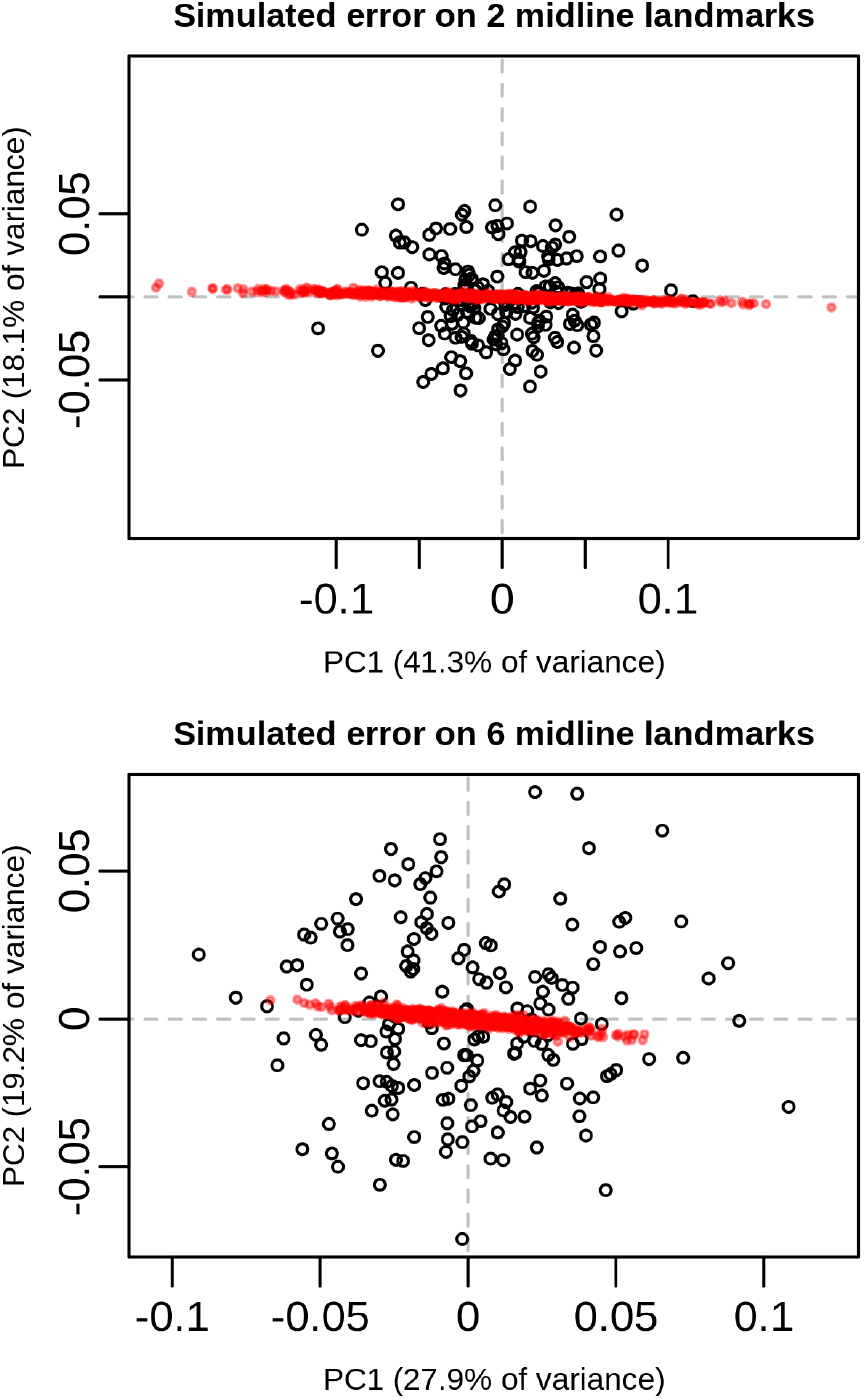
Mirror-reflected simulations on the mean (red) projected into the morphospace of the first two PCs of the symmetrized observations (black). *Top*, 14-landmark arch subset. *Bottom*, 18-landmark full configuration.

The size of a mirror-reflected configuration is similarly dependent on the magnitude of net midline deviations. Figure 8 overlays centroid size densities for the simulations with centroid size densities for the observations. Simulations on the arch subset mean have about half as much centroid size variation as does the entire sample of observations 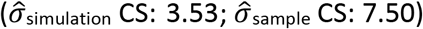. The impact of midline deviations on centroid size is much less for the full configuration 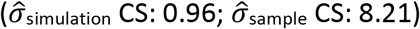.

**Fig 8.**
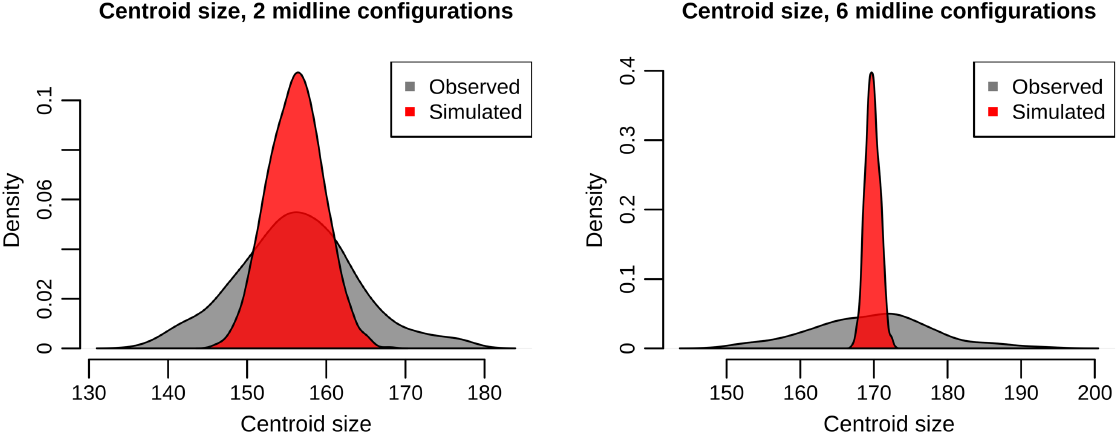
Centroid size variation for observations (black) and simulations of mediolateral error on the mean configuration (red).

### Effect of mirror reflection and shape recovery on sample shape variation

As in Cardini (29, 30), I tested how well samples obtained with several reduced landmarking strategies for efficient data collection approximated patterns of shape variation for complete, bilateral configurations. The shape recovery approach introduced here augments the focal side landmarks with a pair landmarks from the opposite side of the mandible. The criteria for assigning opposite side landmarks to shape recovery pairs was physical distance from the symphysis.

Figure 9 plots the distribution of superimposed shapes for the alternative full configuration data schemes, with mean wireframe superimposed. Relative to mirror reflection, the recovery strategy helps to constrain the effect of midline deviations on shape outcomes. Qualitatively, each of the recovery alternatives looks more similar to the full shape and symmetric superimpositions than does mirror reflection. Recovery with more distal landmarks appears to best replicate bilateral data.

**Fig 9.**
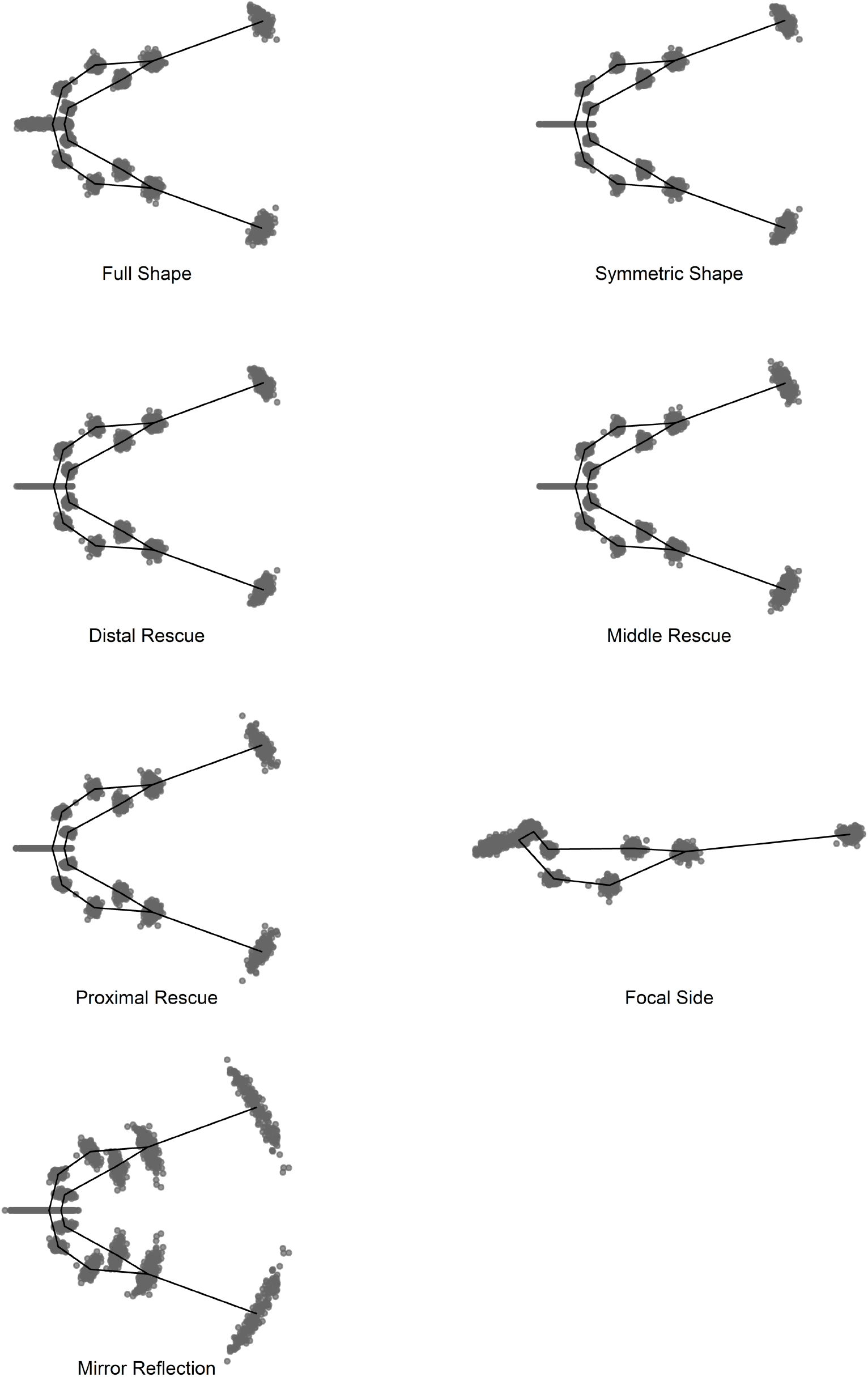
Full configuration superimposed distributions using different data and superimposition (regular vs. symmetric) strategies. Mean wireframe superimposed. The wireframe spans the arch subset landmarks only.

Table 2 provides the pairwise correlations between shape and centroid size distances computed among whole shape (Full), symmetric (Symm.), shape recovery (Distal, Middle, Proximal), focal side (Focal), and mirror-reflected (Mirror) datasets. Supplementary Table 1 provides the same table for the arch subset configurations. For shape (lower triangle), recovery with distal landmarks closely approximates shape variation for datasets that incorporate the full complement of landmarks. Proximal landmark shape recovery and the focal side approach capture shape variation about equally well. The mirror-reflected correlation with the full and symmetric shape distributions is disturbingly weak. For centroid size, all of the efficient data collection strategies replicate full landmark strategies reasonably well (upper triangle), though mirror reflection is clearly the least faithful replica.

**Table 2.** Pairwise shape (lower triangle) and size (upper triangle) correlations between the landmarking and superimposition schemes for full mandible configurations.

|  | Full | Symm. | Shape Rescue |  |  | Focal | Mirror |
| --- | --- | --- | --- | --- | --- | --- | --- |
|  |  |  | Distal | Middle | Proximal |  |  |
| **Full** | 1 | 1 | 0.999 | 0.985 | 0.982 | 0.976 | 0.882 |
| **Symm.** | 0.981 | 1 | 0.999 | 0.985 | 0.982 | 0.976 | 0.882 |
| **Distal** | 0.969 | 0.963 | 1 | 0.979 | 0.976 | 0.973 | 0.883 |
| **Middle** | 0.86 | 0.833 | 0.854 | 1 | 0.991 | 0.984 | 0.846 |
| **Proximal** | 0.678 | 0.643 | 0.712 | 0.724 | 1 | 0.978 | 0.88 |
| **Focal** | 0.727 | 0.696 | 0.738 | 0.697 | 0.584 | 1 | 0.84 |
| **Mirror** | 0.203 | 0.116 | 0.142 | 0.108 | 0.226 | 0.257 | 1 |

## Discussion

Using a simulation approach and comparisons of alternative landmark collection strategies, I evaluated mirror reflection (and a shape recovery alternative) in the context of narrow-midline morphology. Mirror reflection produced highly distorted outcomes. It should be applied with great caution, or not at all, for morphology where the midline is narrow relative to the bilateral landmarks.

In understanding the magnitude of shape and size variation the simulations introduce, it is important to keep in mind that the templates for simulation were means of a diverse human mandible dataset that includes samples from six continents. Despite this global diversity, simulating with a standard deviation of only 0.25 mm produced many observations outside the range of sample variation. Distortions were less severe for the full configuration dataset than the arch subset dataset. The relative improvement in full configuration simulated outcomes is simply due to averaging. As the number of midline landmarks increases from two to six, random deviations from symmetry are more likely to balance out. Moreover, deviations at the mandibular symphysis were mostly expressed along the primary axis of shape variation. This is especially concerning because the mirror reflection effect, while artefactual, looks biologically relevant. It essentially replicates the continuous pattern of shape variation from wider to narrower mandibles that predominates PC 1, and would likely confound statistical analyses and interpretations of shape ordinations.

Shape distortions from mirror reflection produced corresponding, sometimes large, size distortions. As with shape, centroid size was more variable for the arch subset than full configuration simulation. Reduced CS variance in the full configuration simulation has at least two causes. First, more midline landmarks produce a stronger deviation-averaging effect, reducing shape (hence size) distortions. The second reason is more subtle. With more landmarks at the symphysis, the centroid of the full configuration is drawn anteriorly relative to the centroid of the arch subset. For these simulations, which differ primarily in the degree of mandibular flexion from condyle to condyle (see Figure 6), a more anterior centroid will by itself reduce centroid size variation. The **Appendix** provides further discussion of this effect.

Focal side, shape recovery, and mirror reflection alternatives all allow faster data collection than strategies that require the full complement of bilateral landmarks. In the comparisons of these alternatives evaluated herein, mirror reflection provided the least faithful replica of bilateral shape and size data. It is possible that midline measurement error is relatively large for this sample, and that this contributes to the exceptionally poor performance of mirror reflection. Most of the specimens are archaeological. While all are complete cases, many are in imperfect condition. Nevertheless, when the full complement of landmarks is used (Fig. 9 and Table 2 full shape and symmetric superimpositions), mandibular variation for these data appears to be unexceptional. Moreover, the comparatively poor performance of mirror reflection is consistent with the simulation results, and preservation quality is irrelevant to the latter.

The shape recovery approach evaluated herein appears to be a safer strategy for efficient data collection of narrow-midline morphology. Bilateral shape and size relationships were recapitulated very well by incorporating the two most distal landmarks from the opposite side. More proximal opposite side landmarks replicated bilateral configurations less well, but still represented vast improvements over mirror reflection. Ideally, the choice of landmarks would also consider among-landmark variation in measurement error and asymmetry, as these may also influence the quality of a reconstruction. However, the information to do so is lacking for these data.

## Conclusions

Where investigators require large samples and/or many landmarks per specimen, collection of geometric morphometrics data can become a substantial bottleneck in the research process. Efficient data collection strategies can speed a project to completion. In some cases, the improved efficiency may be necessary for a project to be economically or practically feasible. Cardini (29, 30) showed that for some object symmetric morphology, it is possible to obtain reasonably faithful replication of the full complement of bilateral landmarks by collecting landmarks on one side and at the midline, then mirror-reflecting the hemi-configuration to create a complete form.

However, mirror reflection is highly problematic for morphology with a relatively narrow midsagittal. Mirror-reflected outcomes for narrow-midline morphology will be sensitive to measurement error and biological asymmetry at the midline. Morphology with these characteristics include mandibles, some vertebral segments, sacra, and even articulated collections of features (ribcages, pelves).

An alternative strategy augments hemi-form landmarks with a small number of landmarks from the opposite side. However, for some samples, this type of shape recovery is not an option. For archaeological and paleontological data, it is sometimes the case that the available materials include many hemi-samples, preserved only at the midline and on one side. In these cases, a focal side approach is the best alternative. Investigators nevertheless committed to mirror reflection may attempt to visually center the placement of midline landmarks when collecting data, ignoring biological variation. Though this amounts to a decision to deviate from typical landmark definitions, it will likely improve the overall quality of mirror reflection results. In addition, the results presented herein indicate that it is important to have several landmarks at the midline to give any deviations from symmetry the opportunity to average out.

The shape recovery alternative to mirror reflection was able to closely approximate shape and size relationships in the bilateral data. Still, it is important to keep in mind that these improvements are a probability not a promise. Any single reconstructed specimen may substantially deviate from its true bilateral form. Therefore, even the choice to employ a shape recovery strategy requires careful consideration.

---

**Supplementary Table 1.** Pairwise shape (lower triangle) and size (upper triangle) correlations between the landmarking and superimposition schemes for arch subset configurations.

|  | Full | Symm. | Shape Recovery |  |  | Focal | Mirror |
| --- | --- | --- | --- | --- | --- | --- | --- |
|  |  |  | Distal | Middle | Proximal |  |  |
| **Full** | 1 | 1 | 0.996 | 0.981 | 0.97 | 0.953 | -0.033 |
| **Symm.** | 0.975 | 1 | 0.996 | 0.981 | 0.97 | 0.953 | -0.033 |
| **Distal** | 0.962 | 0.95 | 1 | 0.962 | 0.952 | 0.943 | 0.011 |
| **Middle** | 0.772 | 0.716 | 0.771 | 1 | 0.98 | 0.96 | -0.115 |
| **Proximal** | 0.601 | 0.543 | 0.657 | 0.659 | 1 | 0.947 | -0.068 |
| **Focal** | 0.466 | 0.385 | 0.546 | 0.515 | 0.518 | 1 | -0.056 |
| **Mirror** | 0.01 | -0.057 | -0.026 | -0.041 | -0.032 | 0.068 | 1 |

## Appendix Centroid Size Variance as a Function of Centroid Position

An additional reason that centroid size varies less in the full configuration simulation is because, relative to the arch subset, the full configuration centroid is located nearer to the rotational origin of most variation in the simulated dataset, and thus farther from the landmarks that make the largest contribution to shape and size variation. While this demonstration is specific to the type of variation introduced by simulating mediolateral deviations at the symphysis landmarks, it also provides some general intuition about how the spatial distribution of landmarks can influence sample variation.

### Assumptions and simplifications

The working model relies on the following assumptions and simplifications:

1. Variation in the simulated forms is well-approximated by treating the centroid of the symphysis landmarks as the origin of a circle, through which passes a supero-inferior axis of rotation (Figure A1). Simulated observations thus deviate from the mean configuration (simulation template) only in the degree of condyle to condyle flexion or extension. The distribution of simulations depicted in Figure 6 is consistent with this simplification.

**Figure A1.**
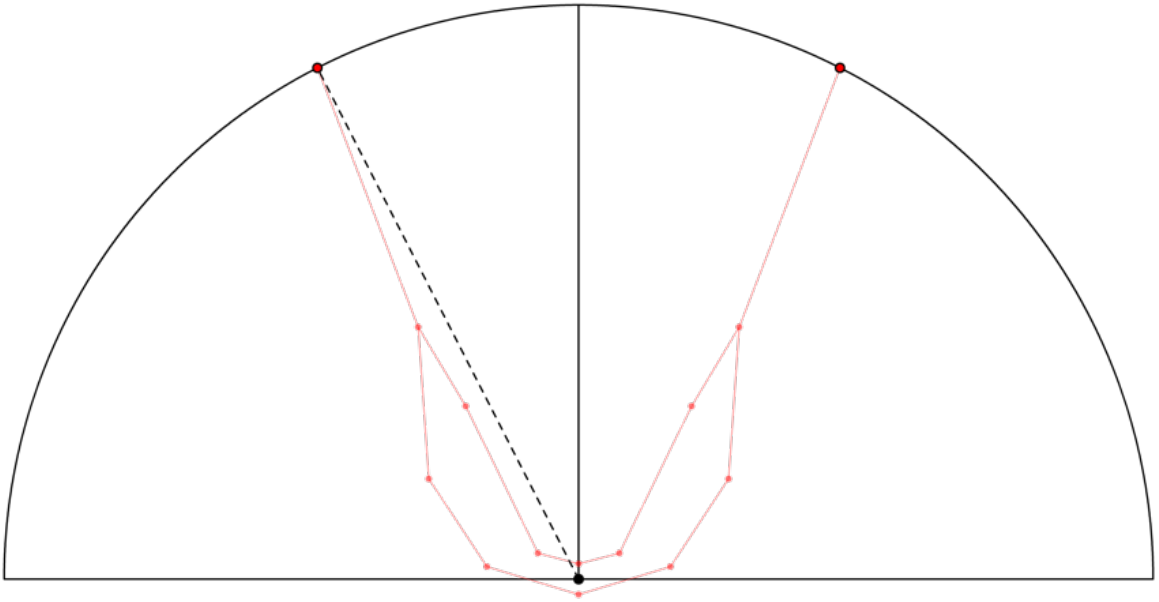
Working model of variation in simulated forms. Red wireframe displays the mean arch subset. The hemi-circle (with origin at black point on symphysis) traces the arc along which the condyle landmarks are constrained to vary. All other bilateral landmarks will vary in unison along smaller radius arcs. From simulation to simulation, the condyles will be closer to or farther from one another, but always because the sides rotate towards (greater flexion) or away from (greater extension) the midsagittal plane, in equal degree about the symphysis centroid.
2. It is clear that condyle position will have the highest variance in the rotational model shown in Figure A1. I therefore simplify the discussion and illustrations by focusing primarily on the relationship between variance in midline rotation and variance in the condyle’s contribution to centroid size variance.
3. The 3D landmark configurations have been reduced to 2D, as shown in Figures A1–A3.
4. I demonstrate the effect on centroid size for differences in condyle position between the mean template and one hypothetical observation. In broad strokes, the implications are easily extended to variation in the centroid size of full configuration, and to a complete sample.

### Centroid position and symphysis landmark count in the arch subset and full configurations

The centroid is the average landmark position for a configuration [1].

The full configuration augments the arch subset with four additional symphysis landmarks. As show in Figure A2, one effect of this augmentation is that the full configuration centroid (open black circle) lies anterior to the arch subset centroid (open red circle).

**Figure A2.**
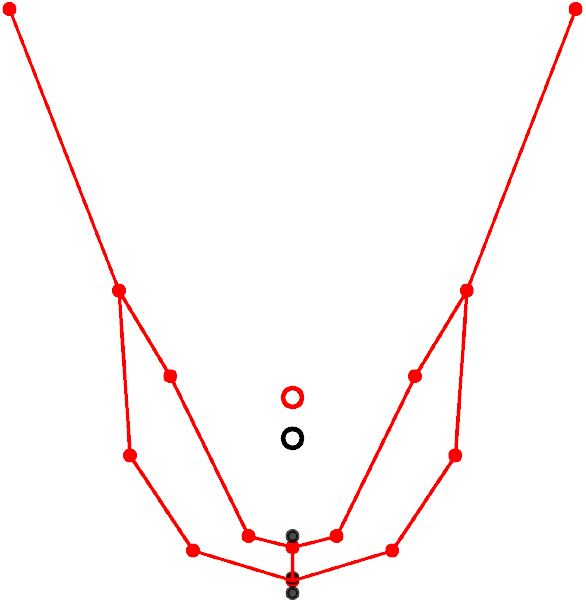
Centroid positions (open circles) for the arch subset (red) and full (black) configurations. The four additional full configuration symphysis landmarks are plotted in black, though only three are visible in this view.

Note that both centroids lie along the midsagittal axis. This will always be the case for the centroid of an object symmetric feature which has been rendered statistically symmetric through reflected relabeling or symmetric superimposition.

### Relationship between centroid size variation and centroid position in configurations that vary by flexion

Centroid size is the square root of the sum of the squared line segment lengths from each landmark to the centroid [1].

In Figure A3, imagine the position of the “centroid” (C*) is permitted to vary freely between the symphysis origin (filled black circle) and some far posterior point along the midsagittal axis. With C* at the origin, no amount of flexion or extension will produce a difference between the line segment lengths from the centroid to the mean and hypothetical right condyle positions (black dashed lines). Thus, midline deviations (mandibular flexion or extension) make no contribution to “centroid size” (CS*) variation. For all C* values posterior to the origin, condyle positions that deviate from the mean condyle position do contribute to variation in CS*, as a function of the difference in lengths from C* to the mean and hypothetical left condyle. These lengths are illustrated for C* on the condylar circumference (filled purple circle; purple dashed lines), and C* where the contribution to CS* variance is greatest, i.e., when C* is positioned such that the line segments to the mean and hypothetical left condyle are collinear (green circle; overlapping green dashed lines).

**Figure A3.**
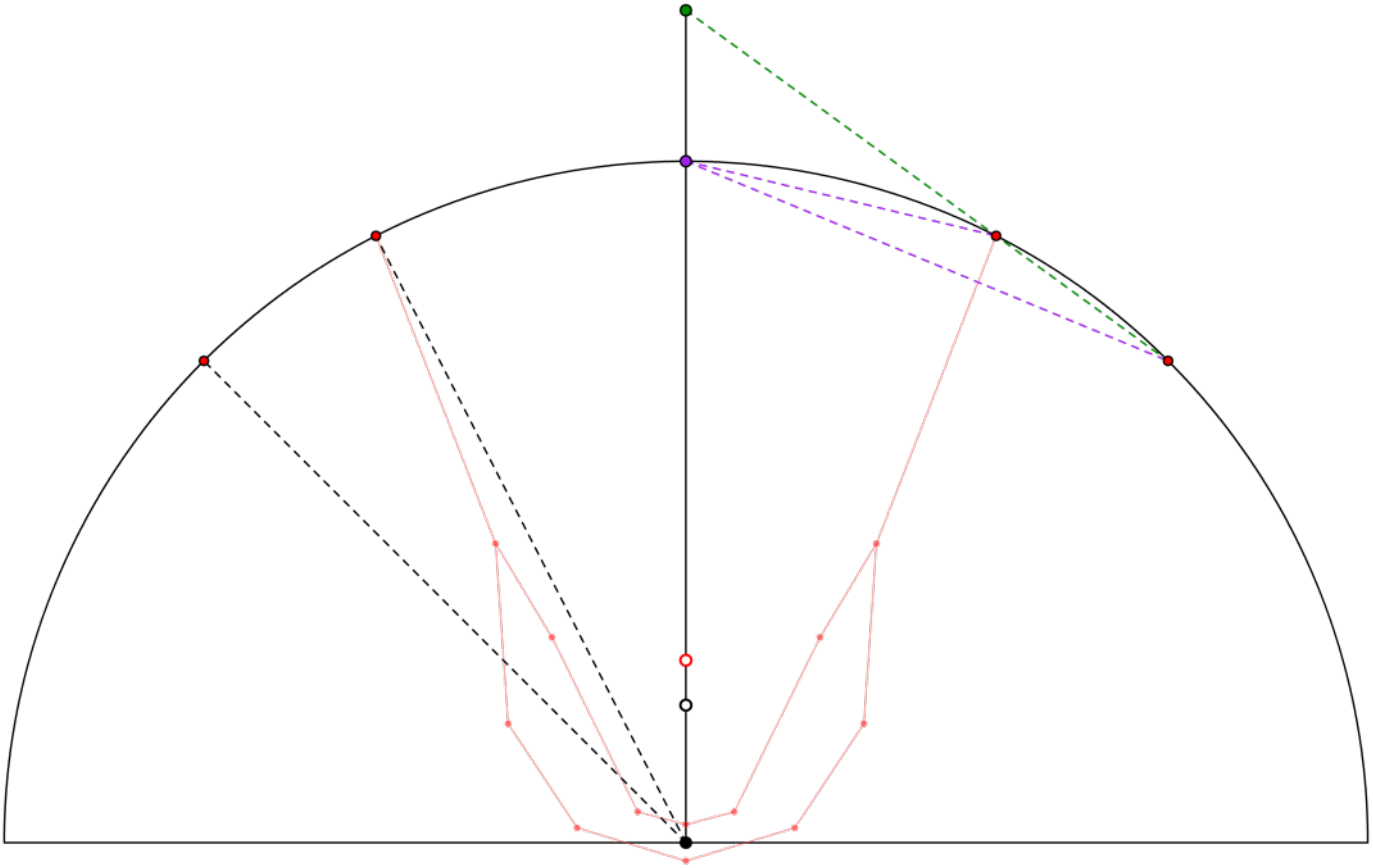
Relationship between C* position and CS* variance contribution. Red landmarks connected by wireframe: mean arch subset. Additional red points along the circumference: hypothetical condyle positions for one observation.

Overall, the contribution of condyle location to CS* variation in this model is a continuous function of the distance of C* from the origin (Figure A4).^1^ At more extreme posterior C* values, the function asymptotically approaches the vertical (y-axis) distance between the mean and alterative condyle positions.

**Figure A4.**
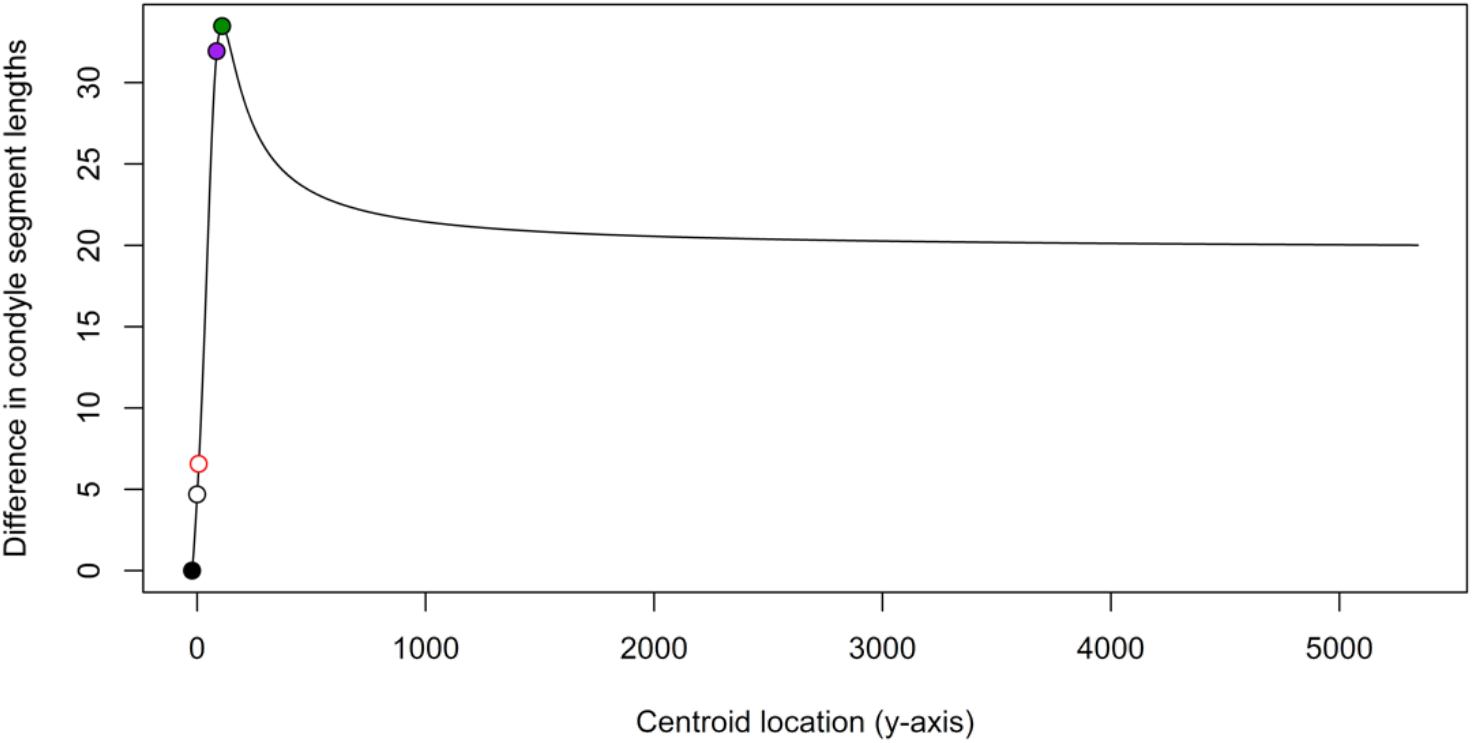
Relationship between C* location and CS* variance contribution for a template condyle and one condyle variant. Closed black circle: variance contribution with C* at origin. Open black circle: with C* at full configuration centroid. Red circle: with C* at arch subset centroid. Purple circle: with C* at intersection of midsagittal radius and circumference. Green circle: C* at maximum chord length difference.

The same effect of C* position on CS* holds when the size differences are computed between a full complement of landmarks for the mean and hypothetical configurations (Figure A5). Whether the mandible is more extended than the mean (solid line and open circles) or more flexed (dashed line and closed circles), a more posterior C* results in greater size differences between the mean and hypothetical configurations.

**Figure A5.**
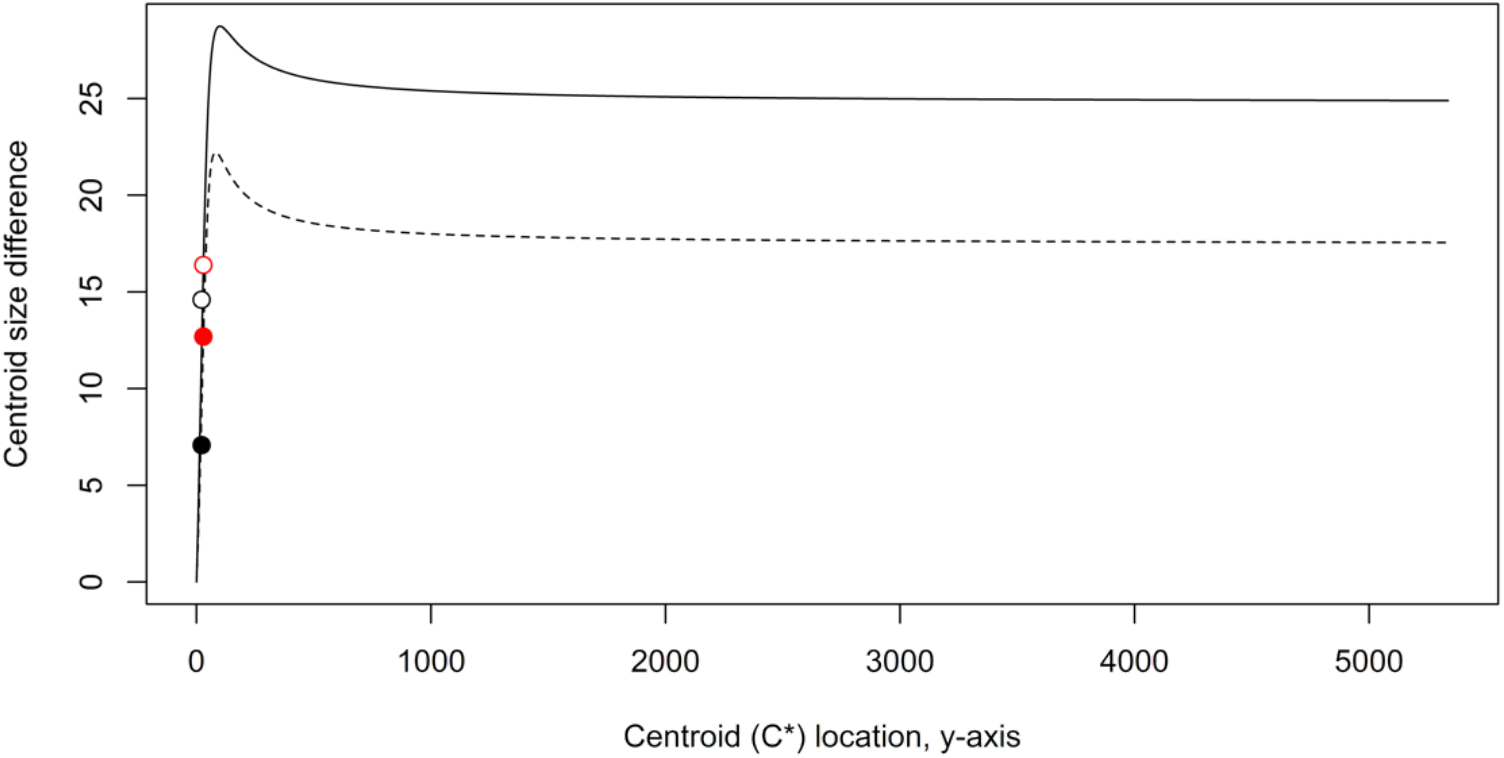
Relationship between C* location and CS* differences between the mean configuration and two simulated configurations. Dashed line with closed circles depicts CS* differences for a configuration that is flexed relative to the mean; solid line with open circles depicts CS* differences for an equally extended configuration. For both configurations, the simulation’s CS* deviation from the mean is lower when C* is derived based on six symphysis landmarks (black points), resulting in a more anterior symphysis, than when it is based on two symphysis landmarks (red points).

In summary, where narrow-midline, symmetric configurations primarily differ from one another by flexion or extension, centroid size variation will depend to some extent on the location of the centroid. Centroid location is in part determined by the landmark map—whether, for instance, many landmarks are concentrated in one region of the morphology of interest, with sparser coverage elsewhere. In our simulations, one reason full configuration size variance is smaller than arch subset size variance is because the full subset has a higher concentration of landmarks at the symphysis. This draws the centroid anteriorly, towards the origin, which ultimately means that, in comparison to the arch subset simulation, variation in bilateral landmark locations in our full configuration simulation has a reduced impact on centroid size.

## Footnotes

1 Extending the model to a sample of condyle positions, condyle position contributions to centroid size variance would be a function of both C* position and the distance of each condyle position from the mean.

## References

1. F. Rohlf, F. Bookstein, Eds., Proceedings of the Michigan Morphometrics Workshop (University of Michigan Museum of Zoology, Ann Arbor, MI, 1988), Vol 2, p 380.

2. F. J. Rohlf, L. F. Marcus, A Revolution in Morphometrics. Trends in Ecology and Evolution 8 (1993).

3. L. F. Marcus, E. Bello, A. Garcia-Valdecasas, Eds., Contributions to Morphometrics (Consejo Superior de Investigaciones Cientificas, Madrid, Spain, 1992), p 264.

4. S. B. Cooke, C. E. Terhune, Form, function, and geometric morphometrics. Anat Rec (Hoboken) 298, 5–28 (2015).

5. D. E. Slice, Geometric morphometrics. Annu. Rev. Anthropol. 36, 261–281 (2007).

6. P. Mitteroecker, F. Bookstein, Linear discrimination, ordination, and the visualization of selection gradients in modern morphometrics. Evolutionary Biology 38, 100–114 (2011).

7. D. C. Katz, M. N. Grote, T. D. Weaver, Changes in human skull morphology across the agricultural transition are consistent with softer diets in preindustrial farming groups. Proc Natl Acad Sci U S A 114, 9050–9055 (2017).

8. C. R. Cooney et al., Mega-evolutionary dynamics of the adaptive radiation of birds. Nature 542, 344–347 (2017).

9. S. R. Frost, L. F. Marcus, F. L. Bookstein, D. P. Reddy, E. Delson, Cranial allometry, phylogeography, and systematics of large-bodied papionins (primates: Cercopithecinae) inferred from geometric morphometric analysis of landmark data. The Anatomical Record Part A: Discoveries in Molecular, Cellular, and Evolutionary Biology 275A, 1048–1072 (2003).

10. M. Pavličev et al., Development Shapes a Consistent Inbreeding Effect in Mouse Crania of Different Line Crosses. J Exp Zool B Mol Dev Evol 326, 474–488 (2016).

11. J. B. Cole et al., Genomewide Association Study of African Children Identifies Association of SCHIP1 and PDE8A with Facial Size and Shape. PLoS Genet 12, e1006174 (2016).

12. M. L. Zelditch, D. L. Swiderski, H. D. Sheets, Geometric Morphometrics for Biologists: A Primer (Elsevier Academic Press, New York, ed. 2nd, 2014).

13. P. Mitteroecker (2018) semilandmarks in biology. in MORPHMET (https://groups.google.com/a/morphometrics.org/forum/#!forum/morphmet).

14. A. Watanabe, How many landmarks are enough to characterize shape and size variation? PLoS One 13, e0198341 (2018).

15. E. Nicholson, K. Harvati, Quantitative analysis of human mandibular shape using three-dimensional geometric morphometrics. American Journal of Physical Anthropology 131, 368–383 (2006).

16. T. Gao, G. S. Yapuncich, I. Daubechies, S. Mukherjee, D. M. Boyer, Development and Assessment of Fully Automated and Globally Transitive Geometric Morphometric Methods, With Application to a Biological Comparative Dataset With High Interspecific Variation. Anat Rec (Hoboken) 301, 636–658 (2018).

17. M. Li et al., Rapid automated landmarking for morphometric analysis of three-dimensional facial scans. J Anat 230, 607–618 (2017).

18. D. M. Boyer et al., Algorithms to automatically quantify the geometric similarity of anatomical surfaces. Proc Natl Acad Sci U S A 108, 18221–18226 (2011).

19. A. M. Maga, N. J. Tustison, B. B. Avants, A population level atlas of Mus musculus craniofacial skeleton and automated image-based shape analysis. J Anat 231, 433–443 (2017).

20. D. Katz, M. Friess, Technical Note: 3D From Standard Digital Photography of Human Crania—A Preliminary Assessment. American journal of physical anthropology 154, 152–158 (2014).

21. C. Klingenberg, Analyzing fluctuating asymmetry with geometric morphometrics: concepts, methods, and applications. Symmetry 7, 843–934 (2015).

22. Y. Savriama, C. Klingenberg, Beyond bilateral symmetry: geometric morphometric methods for any type of symmetry. BMC Evolutionary Biology 11, 280 (2011).

23. K. V. Mardia, F. L. Bookstein, I. J. Moreton, Statistical assessment of bilateral symmetry of shapes. Biometrika 87, 285–300 (2000).

24. C. P. Klingenberg, M. Barluenga, A. Meyer, Shape analysis of symmetric structures: quantifying variation among individuals and asymmetry. Evolution 56, 1909–1920 (2002).

25. K. Harvati, Quantitative analysis of Neanderthal temporal bone morphology using three-dimensional geometric morphometrics. Am J Phys Anthropol 120, 323–338 (2003).

26. P. Gunz, P. Mitteroecker, S. Neubauer, G. W. Weber, F. L. Bookstein, Principles for the virtual reconstruction of hominin crania. Journal of Human Evolution 57, 48–62 (2009).

27. P. Mitteroecker, P. Gunz, Advances in Geometric Morphometrics. Evolutionary Biology 36, 235–247 (2009).

28. F. L. Bookstein (2011) Biology and mathematical imagination: the meaning of morphometrics (Rohlf Medal Lecture).

29. A. Cardini, Lost in the Other Half: Improving Accuracy in Geometric Morphometric Analyses of One Side of Bilaterally Symmetric Structures. Systematic Biology (2016).

30. A. Cardini, Left, right or both? Estimating and improving accuracy of one-side-only geometric morphometric analyses of cranial variation. Journal of Zoological Systematics and Evolutionary Research (2017).

31. A. Cardini, J. Diniz Filho, P. Polly, S. Elton, “Biogeographic analysis using geometric morphometrics: clines in skull size and shape in a widespread African arboreal monkey” in Morphometrics for Nonmorphometricians, A. M. Elewa, Ed. (Springer, 2010), chap. 8, pp. 191–217.

32. F. Bookstein, Size and shape spaces for landmark data in two dimensions. Statistical Science 1, 181–242 (1986).

33. R. FJ, Statistical power comparisons among alternative morphometric methods. American Journal of Physical Anthropology 111, 463–478 (2000).

34. R Development Core Team (2015) R: A language and environment for statistical computing. (R Foundation for Statistical Computing, Vienna, Austria. URL http://www.R-project.org/).

35. D. Adams, M. Collyer, E. Sherratt (2015) geomorph: Software for geometric morphometric analyses. R package version 2.1.6. http://cran.r-project.org/web/packages/geomorph/index.html.

36. I. L. Dryden (2018) shapes package. (R Foundation for Statistical Computing, Vienna, Austria).

37. D. Adler, D. Murdoch, others (2016) rgl: 3D Visualization Using OpenGL.

38. D. E. Slice, “Modern morphometrics” in Modern Morphometrics in Physical Anthropology., D. E. Slice, Ed. (Kluwer Acad./Plenum, New York, 2005), pp. 1–45.

39. B. G, K. R, “Grundlagen der Osteometrie” in Anthropologie. Handbuch der vergleichenden Biologie des Menschen. Band I, 1. Teil., K. R, Ed. (Springer, Stuttgart, 1988), pp. 129–159.

## References

1. Mitteroecker P, Gunz P. Advances in Geometric Morphometrics. Evolutionary Biology. 2009;36(2):235–47. doi: 10.1007/s11692-009-9055-x.

